# Oscillatory waveform shape and temporal spike correlations differ across bat frontal and auditory cortex

**DOI:** 10.1101/2023.07.03.547519

**Authors:** Francisco García-Rosales, Natalie Schaworonkow, Julio C. Hechavarria

## Abstract

Neural oscillations are associated with diverse computations in the mammalian brain. The waveform shape of oscillatory activity measured in cortex relates to local physiology, and can be informative about aberrant or dynamically changing states. However, how waveform shape differs across distant yet functionally and anatomically related cortical regions is largely unknown. In this study, we capitalize on simultaneous recordings of local field potentials (LFPs) in the auditory and frontal cortices of awake, male *Carollia perspicillata* bats to examine, on a cycle-by-cycle basis, waveform shape differences across cortical regions. We find that waveform shape differs markedly in the fronto-auditory circuit even for temporally correlated rhythmic activity in comparable frequency ranges (i.e. in the delta and gamma bands) during spontaneous activity. In addition, we report consistent differences between areas in the variability of waveform shape across individual cycles. A conceptual model predicts higher spike-spike and spike-LFP correlations in regions with more asymmetric shape, a phenomenon that was observed in the data: spike-spike and spike-LFP correlations were higher in frontal cortex. The model suggests a relationship between waveform shape differences and differences in spike correlations across cortical areas. Altogether, these results indicate that oscillatory activity in frontal and auditory cortex possess distinct dynamics related to the anatomical and functional diversity of the fronto-auditory circuit.

**Significance statement:** The brain activity of many animals displays intricate oscillations, which are usually characterized in terms of their frequency and amplitude. Here, we study oscillations from the bat frontal and auditory cortices on a cycle-by-cycle basis, additionally focusing on their characteristic waveform shape. The study reveals clear differences across regions in waveform shape and oscillatory regularity, even when the frequency of the oscillations is similar. A conceptual model predicts that more asymmetric waveforms result from stronger correlations between neural spikes and electrical field activity. Such predictions were supported by the data. The findings shed light onto the unique properties of different cortical areas, providing key insights into the distinctive physiology and functional diversity within the fronto-auditory circuit.

## Introduction

Rhythmic neural activity at various timescales underpins several functions in the mammalian brain. In the frontal cortex, oscillations of local-field potentials (LFPs) in low and high frequencies are implicated in cognitive and executive control (Helfrich and Knight, 2019; Insel et al., 2012; Rajan et al., 2019; Tavares and Tort, 2022; Veniero et al., 2021; Zhang et al., 2016), while rhythmic activity in sensory cortices is linked with the effective encoding of incoming stimuli (Gourevitch et al., 2020; Gross et al., 2007; Kienitz et al., 2021; Lakatos et al., 2007; Tan et al., 2019; Teng et al., 2017; Uran et al., 2022). These oscillations reflect the underlying dynamics of their generating motifs, which determine several of their properties, including waveform shape (Cole and Voytek, 2017). Indeed, waveform shape and related features change in the developing brain (Schaworonkow and Voytek, 2021) and possess atypical characteristics in disease (Cole and Voytek, 2019; Cole et al., 2017; Jackson et al., 2019). Waveform patterns of oscillatory activity can provide important insights into the physiology and function of the neocortex, yet how they differ across cortical regions remains largely unstudied.

In this work, we examine oscillatory waveform shape in the frontal and auditory cortices of a mammalian vocal specialist, the bat *Carollia perspicillata*. The bat auditory cortex (AC) is a well-studied structure that presents both spontaneous and stimulus-driven rhythmic patterns of neuronal activity (Garcia-Rosales et al., 2019; Hechavarria et al., 2016; Medvedev and Kanwal, 2004). As in other mammals (Lakatos et al., 2005; Luo and Poeppel, 2007; Neymotin et al., 2022; Teng et al., 2017), LFPs in the bat AC track the temporal dynamics of acoustic sequences with periodic and quasi-periodic temporal structures (Garcia-Rosales et al., 2018). LFPs in *C. perspicillata*’s AC exhibit clear coupling with neuronal spiking, potentially coordinating single-cell responses to acoustic stimuli and contributing actively to the encoding of multi-syllabic communication sounds (Garcia-Rosales et al., 2018).

In the frontal cortex, we focused on the frontal auditory field (FAF), a structure specialized in auditory-related behaviour (Eiermann and Esser, 2000; Kanwal et al., 2000; Kobler et al., 1987). This region is anatomically connected with the AC, but receives also relatively fast inputs from an alternative pathway bypassing midbrain and cortex (Kobler et al., 1987). Pre- and post-vocal dynamics in the FAF, as well as its functional connectivity patterns with the AC and the striatum, implicate this region in the control of vocalization behaviour (Garcia-Rosales et al., 2022b; Weineck et al., 2020). Furthermore, the FAF is anatomically connected with the superior colliculus, suggesting that it may be involved in coordinating fast movements based on the bat’s auditory environment (Casseday et al., 1989; Kobler et al., 1987). The nature of FAF-AC interconnectivity suggests that the FAF plays a crucial role in the integration of auditory feedback for the coordination of rapid auditory-based behaviour (Garcia-Rosales et al., 2022b). These data indicate that, while the AC operates as a classical sensory cortex, the FAF acts as part of a control and integration hub.

While low- and high-frequency oscillatory activities in the bat FAF-AC network are functionally related, it is unknown how they differ in terms of waveform shape. Characterizing waveform shape differences across cortical regions could be an informative step towards understanding how neuronal oscillations in these areas differ, and thus constrain hypotheses about the mechanisms underlying neural activity across structures. By means of simultaneous electrophysiological recordings and cycle-by-cycle analyses of neural oscillations, we show that the waveform shape and variability of frontal- and auditory-cortical oscillations differ markedly in delta and gamma frequencies. We demonstrate a relationship between waveform shape and spike correlations by modelling and computing spike-field measures. We argue that these differences reflect physiological disparities in the FAF-AC circuit, and establish a potential link between spike timing and waveform shape. Our results support the notion of heterogeneity of cortical rhythms in the mammalian brain, and stress the importance of waveform shape for understanding cortical physiology and function.

## Materials and Methods

### Animal preparation and surgical procedures

The study was conducted on two awake *Carollia perspicillata* bats (2 males), which were obtained from a colony at the Goethe University, Frankfurt. All experimental procedures were in compliance with European regulation and were approved by the relevant local authorities (Regierungspräsidium Darmstadt, experimental permit #FU-1126). Animals used in experiments were kept isolated from the main colony, with a reversed light-dark cycle (i.e. lights off from 12:00 to 00:00; this applies to all bats in the colony as well).

The data presented in this work were collected as part of a previous study (Garcia-Rosales et al., 2022b), where a detailed description of the surgical procedures can be found. In brief, bats were anesthetized with a mixture of ketamine (10 mg*kg^−1^, Ketavet, Pfizer) and xylazine (38 mg*kg^−1^, Rompun, Bayer), and underwent surgery in order to expose the skull in the areas of the frontal and auditory cortices. A metal rod (ca. 1 cm length, 0.1 cm diameter) was glued onto the bone for head fixation during electrophysiological recordings. A local anaesthetic (ropivacaine hydrochloride, 2 mg/ml, Fresenius Kabi, Germany) was applied subcutaneously around the scalp area prior any handling of the wounds. The precise locations of the FAF and AC were determined by means of well-described landmarks, including the sulcus anterior and prominent blood vessel patterns (Eiermann and Esser, 2000; Esser and Eiermann, 1999; Garcia-Rosales et al., 2020). Access to the frontal and auditory regions of the left hemisphere was gained by cutting small holes (ca. 1 mm^2^) with a scalpel blade on the first day of recordings. Electrophysiological recordings in the AC were made mostly in the high frequency fields (Esser and Eiermann, 1999).

After the surgery animals were given sufficient time to recover (no less than 2 days) before the beginning of experimental sessions. A session did not last more than 4 hours per day. Water was offered to the bats every 1 – 1.5 hours. Experiments were halted if an animal showed any signs of discomfort (e.g. as excessive movement). No animal was used on two consecutive days for recordings.

### Electrophysiological recordings

Electrophysiological measurements were made acutely from fully awake animals in a sound-proofed and electrically isolated chamber. Inside the chamber, bats were placed on a custom-made holder kept at a constant temperature of 30 °C using a heating blanket (Harvard, Homeothermic blanket control unit). Data were acquired simultaneously from the FAF and AC of the left hemisphere using two 16-channel laminar probes (Model A1×16, NeuroNexus, MI; 50 μm channel spacing, impedance: 0.5–3 MΩ per electrode). For each paired FAF-AC recording, probes were carefully inserted into the tissue using piezo manipulators (one per probe; PM-101, Science products GmbH, Hofheim, Germany), perpendicular to the cortical surface, until the top channel was barely visible above the surface. The typical width of *C. perspicillata*’s cortex, and the total span of the electrodes in the probes (750 μm) allowed us to record from all six cortical layers at once (see Garcia-Rosales et al. (2022b); Garcia-Rosales et al. (2019)). From one paired recording to the next in the same experimental session, probes were retracted from the cortical tissue and moved to a new location within the craniotomy in FAF or AC, as distant as possible form previous recording sites within that craniotomy. New recording locations were chosen at the beginning of each experimental session.

Probes in FAF and AC were connected to micro-preamplifiers (MPA 16, Multichannel Systems MCS GmbH, Reutlingen, Germany), while acquisition was done with a single 32-channel system with integrated digitization (sampling frequency, 20 kHz; precision, 16 bits) and amplification steps (Multi Channel Systems MCS GmbH, model ME32 System, Germany). Silver wires were used as references electrodes for each recording shank (i.e. in FAF and AC) placed at different areas of the brain (for FAF: non-auditory lateral ipsilateral region; for AC: non-auditory occipital ipsilateral region). The silver wires were carefully positioned between the skull and the dura matter. The reference and the ground of each probe were short-circuited, and the ground was ultimately common in the acquisition system (the ME32). Recordings were monitored online and stored in a computer using the MC_Rack_Software (Multi Channel Systems MCS GmbH, Reutlingen, Germany; version 4.6.2). Due to technical reasons, the signal from one FAF channel (depth: 500 μm) was linearly interpolated from its immediate neighbours.

### Pre-processing of spiking and LFP signals

All data analyses were made using custom-written Python scripts. Raw data from the recording system were converted to H5 format using Multichannel System’s *McsPyDataTools* package (https://github.com/multichannelsystems/McsPyDataTools, version 0.4.3), and were then parsed and handled with *Syncopy* (https://github.com/esi-neuroscience/syncopy, version 2022.8). Local-field potentials were obtained by filtering the raw data with a low pass Butterworth filter (4^th^ order) with a cut-off frequency of 300 Hz. For computational convenience, LFP signals were then downsampled to 5 kHz. On occasions a sharp spectral peak at 100 Hz was present in the recordings, corresponding to a harmonic of the line noise. We were discouraged to use LFPs close (or above) to 100 Hz in the analyses for the following reasons: (i) frequencies close to 100 Hz would be affected by the line noise harmonic; and (ii) high frequency LFPs (> 100 Hz) can be directly influenced by spiking activity in the form of, for example, spike-bleed through (Ray, 2015). Spike bleed-through constitutes a potential confound that we sought to avoid.

For the detection of multi-unit activity, the raw data was bandpass filtered between 300 and 3000 Hz with a 4^th^ order Butterworth filter. Spikes were detected based on threshold crossing: we defined a spike as a peak with an amplitude of at least 3.5 standard deviations relative all samples in the signal. Only peaks separated by at least 2 ms were considered.

### Spectral analyses

Power spectral densities (PSDs) were computed using Welch’s method (segment length 20480 samples, i.e. 4096 ms) implemented in *scipy* (version 1.9.1). PSDs were calculated independently for each LFP trace (all channels in the N = 29 recordings in both FAF and AC). LFP traces were typically *circa*, but not shorter than, 1200 s long (median: 1239.5 s; 25^th^ percentile: 1252.8 s; 75^th^ percentile: 1423.9 s). The power of each recording was parametrized using a spectral parametrization model (Donoghue et al., 2020), with which a 1/f fit of the PSD was computed. All fits had an *R*^2^ > 0.93 (mean: 0.9965, s.e.m.: 0.001).

We reasoned that significant deviations of the power spectrum from the 1/f fit potentially represented oscillatory activity at a given frequency range. Thus, we normalized each power spectrum by its 1/f component to highlight spectral peaks in FAF and AC. Normalized values would hover around 0 in the case of no spectral peaks, and would be consistently greater than 0 for frequencies in which LFPs presented clear deviations from the underlying 1/f trend. For each channel, we considered a significant deviation from the 1/f if the normalized power at a certain frequency was significantly larger than 0 (FDR-corrected, two-sided one-sample t-tests, p_corr_ < 0.05). This analysis was done for each individual animal (Bat-01, N = 15; Bat-02, N = 14), for frequencies ranging from 1 to 120 Hz. From the results in the two bats, we established the following frequency bands of interest: delta (1–4 Hz) and gamma (65–85 Hz).

### Cycle-by-cycle analyses

For detecting oscillatory bursts in the frequencies of interest we used the *bycycle* package ((Cole and Voytek, 2019), version 1.0.0). The *bycycle* algorithm makes it possible to detect individual cycles in frequency range of interest (here, the frequency bands outlined above), and then to determine whether detected cycles belong to so-called “oscillatory bursts”. An oscillatory burst consists of a sequence of cycles (at least 3 in this study) with stable temporal properties that are mainly summarized as follows: amplitude consistency, period consistency, and monotonicity (rise and decay flanks of cycles in a burst should be mostly monotonic). Furthermore, one parameter controls for signal-to-noise ratio (SNR): the amplitude fraction threshold (see **Fig. 2A**). This parameter rejects cycles whose amplitudes are below a certain percentile relative to the amplitude of all cycles in a given trace. As in (Schaworonkow and Voytek, 2021), we chose the following thresholds for cycle detection: Amplitude fraction threshold, 0.5; Amplitude consistency threshold, 0.5; Period consistency threshold, 0.5; Monotonicity threshold, 0.5.

Each cycle was characterized according to the following features, which determine waveform shape: cycle period (i.e. the duration of each cycle), cycle rise-decay asymmetry (the asymmetry between rise and decay times in the cycle), and cycle peak-trough asymmetry (the asymmetry in duty cycle; see also (Cole and Voytek, 2019; Schaworonkow and Voytek, 2021)). Bursts were characterized according to their duration (the sum of the individual duration of each cycle in the burst). Only cycles that were part of oscillatory bursts were used for further analyses.

To compare cycle features across different recording sites (e.g. between channels in FAF and AC), we defined the value of a given feature for a certain recording as the median value of that feature across all detected cycles in the recording. This was made per LFP trace, therefore yielding one value per recording site (N = 29 paired FAF-AC sites, across 16 channels; data from the two bats were pooled as spectral and bursting patterns were highly consistent across animals). Given that data from FAF and AC were acquired simultaneously for each paired recording, the above allowed us to compare across sites using paired statistics (FDR-corrected Wilcoxon signed-rank tests, significant for p_corr_ < 0.05). Only values derived from simultaneously recorded LFP traces were compared to one another.

A median asymmetry value of 0.5 for a given LFP trace indicates that cycles tend towards a sinusoidal shape. The farther the value is from 0.5 (above or below) the more asymmetric a waveform is. However, whether such values lie above or below the 0.5 threshold strongly depends on signal polarity. Note, for example, that a certain signal and its copy, the latter with inverse polarity, will have values of asymmetry equally distanced from 0.5, but in opposite directions (as peaks become troughs with a polarity inversion). Thus, not controlling for signal polarity can be a strong confound when comparing waveform shape asymmetries, especially if these are calculated from electrodes located in different brain regions which already have dissimilar cytoarchitectures, such as the frontal and auditory cortices. Since we are unable to control for signal polarity in the current dataset, we avoid this potential confound by normalizing median asymmetry values to 0.5. That is, the asymmetry value for a given LFP trace used for comparisons is given by the absolute value of the difference between its raw asymmetry and 0.5. This approach measures how far from sinusoidal the waveform shape of an LFP trace is, independently of signal polarity (Schaworonkow and Voytek, 2021), and is therefore better suited for inter-areal comparisons of waveform shape asymmetry.

The aforementioned cycle features characterize waveform shape, but they do not quantify to what degree individual burst cycles in a given LFP trace are similar to one another. This is measured by the dispersion of the distribution of the cycle features, which was quantified here as the coefficient of variation (CV). The CV is computed over each LFP trace, therefore quantifying cycle-by-cycle the variability over time; it is expressed as follows:

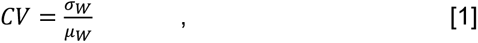

where σ_*W*_ is the standard deviation of the cycle feature distribution (*W*), and μ_*W*_ its mean.

Every recording in frontal or auditory cortex had a specific CV for a given cycle parameter, channel and frequency band. As described with median feature values, this enabled us to conduct paired statistics when comparing CV values between FAF and AC (FDR-corrected Wilcoxon signed-rank tests, significant for p_corr_ < 0.05). The CV was calculated from raw feature values (not normalized to 0.5) across all cycles in a given signal, given that this metric is not affected by signal polarity as it is self-contained for each LFP trace. This allows to explore cycle-by-cycle variability over time without affecting the individual asymmetry values of the cycles involved.

### Sensitivity analyses

To evaluate the dependence of significant differences across cortical structures on the burst detection parameters of the *bycycle* algorithm, bursts were detected as above but detection parameters were varied in pairs as follows: (i) amplitude fraction threshold vs. amplitude consistency threshold; (ii) amplitude fraction threshold vs. period consistency threshold; and (iii) amplitude fraction threshold vs. monotonicity threshold. The same parameter values were used to detect bursts in FAF and AC. However, we also evaluated to what degree our results were sensitive to different burst detection parameters across regions, varying the amplitude fraction threshold independently in each area (**Fig. 7**). Parameters were varied in the range from 0.1 to 0.9, with a step of 0.1. All waveform features were computed as described above, and the variability of waveform features was measured as the CV. As in the original analyses, all channels were statistically compared against each other. We then determined the median of the effect size of comparisons across areas (i.e. the median effect size of the upper-right quadrant of the comparison matrices in **Figs. 5, 6**; effect sizes of non-significant comparisons were set to 0), and plotted this median against parameter combinations (**Figs. 7**) to determine how changing detection parameters affected the reported inter-areal differences.

### Burst co-occurrence analysis

The relationship between the onset of a burst in a specific channel and the cumulative burst co-occurrence with all other channels was calculated as follows. First, given a certain channel (e.g. channel A, for convenience) we determined the onset of all bursts detected across all recordings (the data from the two bats were pooled given that spectral and bursting patterns were highly consistent across animals). In a time window centred on each burst (spanning from -1000 to 1000 ms for delta frequencies, and from -250 to 250 ms for the gamma frequencies) we counted and accumulated, for every channel, the time points at which bursts occurred. Bursts were counted even if their onset or offset were outside the aforementioned window, as long as at least a segment of the burst occurred within that window. For every given channel (channel A in this example) this procedure yielded a matrix (dimensions: [channels x samples]) whose values indicate the accumulated, co-occurring bursting activity in each other electrode, relative to the times in which a burst onset occurred in the channel of interest (i.e. A in this case). We referred to this matrix as a channel’s burst co-occurrence matrix. Burst co-occurrence matrices were computed for 8 channels, 4 in the FAF and 4 in the AC (at depths of 50, 250, 450, and 700 μm). This reduced computational costs and facilitated visualization, while at the same time allowing to explore burst co-occurrences at various depths in each region including superficial, middle and deep layers of cortex.

In order to evaluate whether the onset of a burst in a given electrode was related to the occurrence of bursts in other electrodes, the burst co-occurrence matrix for the channel of interest (e.g. channel A) was normalized following a bootstrapping procedure. We calculated 500 bootstrap burst co-occurrence matrices, but instead of utilizing burst onsets as a reference, pseudo-random time points were used. Because the accumulated number of co-occurring bursts across channels depends on the number of burst onsets used from the reference channel (A), we ensured that the number of randomized time points was equal to the number of burst onsets individually for each recording. The 500 pseudo-random burst co-occurrence matrices were used as a baseline distribution, and the values of the burst co-occurrence matrix for the channel of interest A were then Z-normalized relative to the bootstrap matrices. Absolute Z-score values ≥ 6 were considered significant. Note that the Bonferroni correction of an alpha of 0.05 over 32 channels, 2 frequency bands, 8 channels of interest and 1000 time points yields a significance threshold of 9.7×10^-8^, equivalent to a Z-score of 5.2. In our analysis, negative Z-scores indicate a suppression of burst activity relative to baseline, while positive values indicate an enhancement. These procedures are illustrated in **Fig. 4A**.

### A conceptual model of spike correlations and LFP waveform shape

Synthetic spike trains were modelled as inhomogeneous Poisson processes with firing rates controlled by a pulse train with a frequency of 3 Hz. The duty cycle of the pulse train defines a temporal window in which spiking occurs. Narrow spiking windows (i.e. lower duty cycles) constrain the firing of a neural population in time, resulting in increased temporal correlations across neurons. By contrast, wider spiking windows (i.e. higher duty cycles) result in decreased temporal spiking correlations across neurons. By systematically adjusting the duty cycles we were therefore able to explore how temporal correlations in a neuronal population (N = 30 neurons in our model) might affect LFP waveform shape.

From the spiking activity we derived synthetic LFP signals as follows. The spike train of each simulated neuron was convoluted with a synaptic kernel whose rise and decay times were set to 1 and 20 ms, respectively (function *sim_synaptic_kernel* of the python package NeuroDSP, version 2.2.1; (Cole et al., 2019)). The sum of all convolutions was taken as the LFP signal. The procedure is illustrated in **Fig. 8A**. We generated 300 s of spikes and LFPs for several values of duty cycles (5% to 60%, step: 5%). Cycle features were extracted from the synthetic LFPs by applying of the *bycycle* algorithm described above.

### Spike-spike correlations

All detected spiking events (see above) were included to calculate spike train correlations across channels. Spike trains were binned using 5 ms bins, and the Pearson’s correlation coefficient across pairs of binned spike trains was computed using the *Elephant* toolbox (v. 0.12.0; https://github.com/NeuralEnsemble/elephant). Correlation coefficients from channels located in the FAF were averaged, and the same was done for channels located in the AC. This yielded one correlation value per recording in FAF and AC, which allowed to capitalize on simultaneous recordings in both regions by means of paired statistical comparisons (Wilcoxon signed-rank test, alpha = 0.05).

### Pairwise phase consistency

The pairwise phase consistency (PPC) was computed as described in previous literature (Vinck et al., 2010). Only spikes that occurred within oscillatory bursts in FAF or AC were considered. If more than 10000 spikes were detected in a given trace, 10000 spikes were randomly selected to calculate PPC given that analyses were computationally expensive for large spike counts. In order to minimize the risk of asymmetric signals yielding unclear measurements of phase, spike phases were not obtained by means of a Hilbert transform or a Fourier analysis. Instead, the timing of a spike was expressed as the time in which the event occurred relative the onset and offset of a cycle as detected in the time series by the *bycycle* algorithm. Thus, each spike timing was between 0 and 1 (0 being the beginning of a burst cycle, 1 being the end), and was converted to a phase by multiplication with 2π. These phases were then used for PPC calculation, which can be expressed as follows (Vinck et al., 2010):

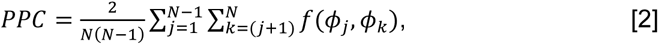

where N is the number of spikes, and ϕ_*j*_, ϕ_*k*_ represent the phases of spikes *j* and *k*, respectively. The function *f*(ϕ_*j*_, ϕ_*k*_) calculates the dot product between two unit vectors. It can be expressed as follows:

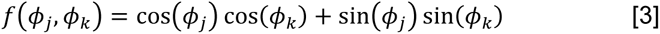

PPC values were averaged in FAF and AC, and paired statistical comparisons were made to evaluate whether significant differences in spike phase consistency existed between regions (Wilcoxon signed-rank test, alpha = 0.05).

### Statistical analyses

All statistical analyses were performed using *scipy* (version 1.9.1), or custom written Python scripts. For determining significant deviations from a 1/f fitted trend in the LFP spectra one-sample t-tests were performed. Statistical comparisons of median and CV values across regions (and within regions) were performed using paired statistics (Wilcoxon signed-rank tests, alpha = 0.05), as recordings in FAF and AC were performed simultaneously (N = 29). Comparisons of spike-spike and spike-LFP correlation (PPC values) were also made using paired statistics. Comparisons of burst lengths were made by means of non-paired statistics (Wilcoxon rank-sum tests, alpha = 0.05). All tests were corrected for multiple comparisons using the false discovery rate when appropriate (Benjaminin and Hochberg procedure (Benjamini and Hochberg, 1995)); it is noted in the main text whenever this correction was applied.

## Results

### Spectral properties of frontal and auditory cortical LFPs

A total of 29 paired recordings (i.e. simultaneous electrophysiological acquisition from each region) in FAF and AC were performed in two bats: Bat-01 and Bat-02 (N = 15 and N = 14 paired FAF-AC recordings, respectively). A schematic representation of the laminar probes, channel depths, and recording locations in the AC are given in **Fig. 1A**. Since a clear map of the FAF does not exist, it was not possible to map electrode locations in the frontal structure in a similar manner. Example frontal and auditory cortical LFP traces from both animals are given in **Fig. 1B, K**, across all recording depths. Typically, LFPs exhibited clear rhythmicity in low and high frequencies in both cortical regions. Evidence for rhythmic activity was clear in grand-average spectra obtained from all ∼20-min long LFP traces (**Fig. 1C, G, L, M**). Observable “bumps” in these spectra are interpretable as deviations from a 1/f power-law (a property of aperiodic mesoscopic signals such as LFPs (Baranauskas et al., 2012)) and therefore suggest the presence of oscillatory activity. We performed spectral parametrization by fitting 1/f curves to the power spectral density of every LFP signal recorded (Donoghue et al., 2020) in order to confirm that such spectral bumps were in fact significant deviations from an aperiodic spectrum. Representative parametrized spectra are depicted in **Fig. 1D, H, M, Q**, corresponding to the full ∼20-min LFP traces from which data in **Fig. 1B, K** were selected. The 1/f fit is shown in dashed blue lines. Deviations in the spectra from the power-law trend were clear in both animals, particularly in the FAF. We tested whether such deviations were consistently present in all recordings by normalizing the power spectrum of each LFP trace (N = 15 in Bat-01, N = 14 in Bat-02, per channel) to their fitted 1/f function (**Fig. 1E, I, N, R**). Power spectral values would hover around 0 in the absence of consistent deviations, but would be significantly above zero otherwise. Normalized spectral values were significantly above 0 in FAF and AC for both animals (FDR-corrected one-sample t-tests; p_corr_ < 1.73×10^-4^, t > 2.25) at relatively low (∼1–5 Hz in FAF and AC), intermediate (∼12–27 Hz in AC), and relatively high (ranging from ∼32–105 Hz, but peaking at 70-85 Hz in FAF and AC) frequencies (**Fig. 1F, J, O, S**).

**Figure 1.**
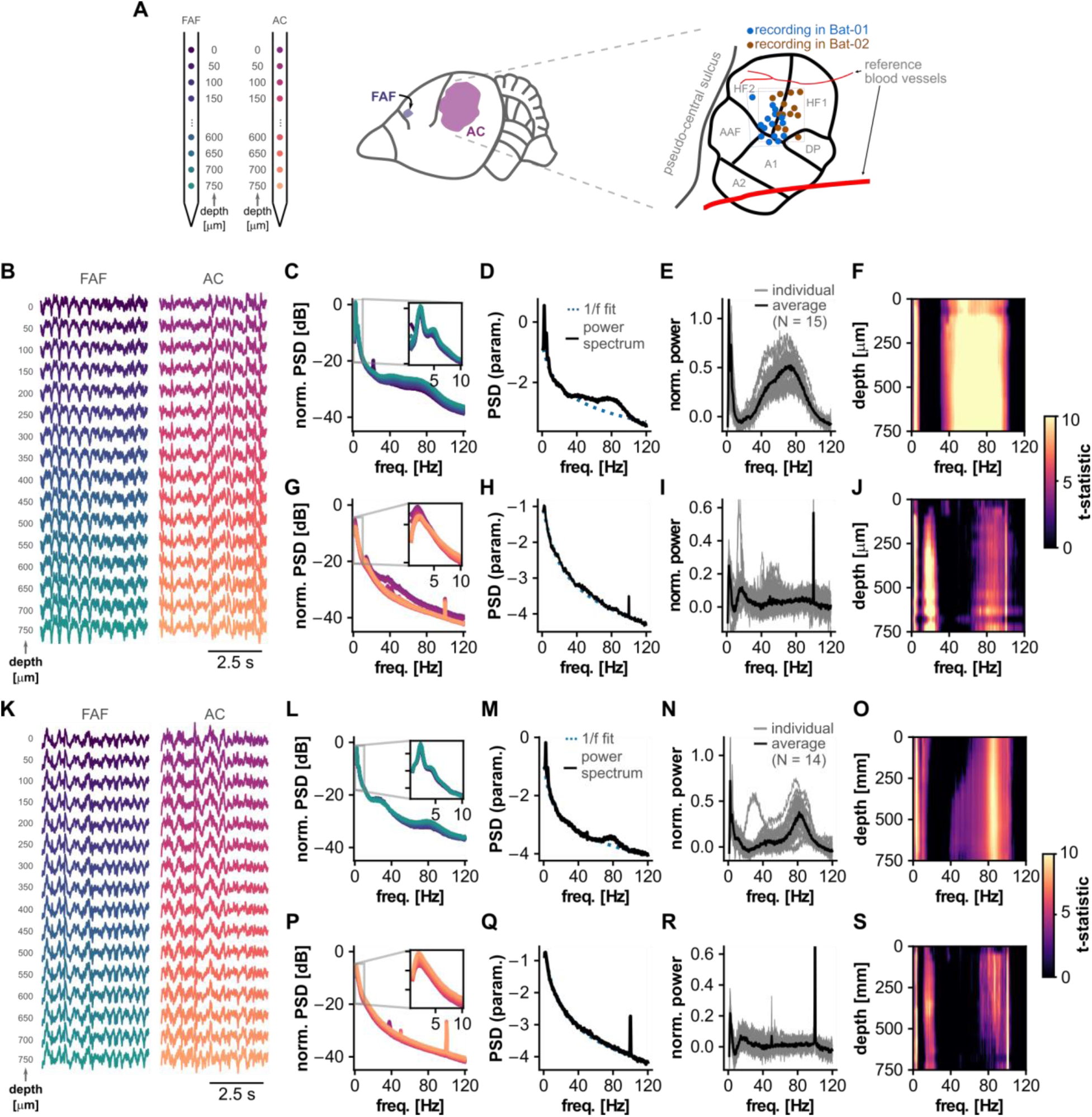
Spectral properties of local-field potentials in FAF and AC. (**A**) *Left*: Schematic representation of the probes used for recordings in FAF and AC. Depth and channel colours correspond to those in panels **B** and **K**. *Middle*: Location of the FAF and AC in *C. perspicillata*’s cortex. *Right*: Schematic representation of recording locations in AC, colour-coded by animal (blue: Bat-01, N = 15; brown: Bat-02, N = 14). The precise location of one recording in Bat-02 could not be recovered. (**B**) Cortical LFPs (5 s excerpts) recorded simultaneously from the FAF (left) and AC (right; note that channel depths are colour-coded) of Bat-01. (**C**) Average power spectra in FAF across all recordings (N = 15) in Bat-01 using full LFP traces (lengths of ∼20 minutes), for all channels. The spectrum of each channel is colour-coded according to the depth scheme of panels **A**, **B**. (**D**) Parametrization of an exemplary power spectrum obtained from ∼20 minutes LFP recordings in the FAF (depth, 700 μm). LFP traces originate from the same recording shown in **B**. The 1/f fit is depicted as a blue dashed line; power spectrum shown in solid black. (**E**) Normalized power spectra (to 1/f activity) across all recordings in Bat-01, shown for channels located at 700 μm in FAF. Solid black line indicates average (N = 15). (**F**) We tested whether the normalized power spectrum was significantly larger than 0 (FDR-corrected t-test, pcorr < 0.05) across depths and frequencies. The t-statistics are summarized here; values were set to 0 if the normalized power spectrum was not significantly (i.e. pcorr >= 0.05). (**G-J**) Similar to panels **C**-**F**, but data shown corresponds to channels located in the AC. (**K**-**S**) Same as **B**-**J** but with data recorded from Bat-02.

In bats such as *C. perspicillata* (the animal studied here), LFP activity in low and high frequencies is related to vocal production (e.g. at frequencies delta: 1-4 Hz, beta: 12–30 Hz, and gamma: 60–120 Hz; see Garcia-Rosales et al. (2022b); Weineck et al. (2020)). Considering the above and the patterns of deviations from a pure 1/f signal shown in **Fig. 1**, in subsequent analyses we focused on frequency bands delta (1–4 Hz) and gamma (65–85 Hz). Beta-band frequencies were not included because no clear peaks in this range were detected in FAF signals (**Fig. 1F, J, O, S**).

### Cycle-by-cycle analysis of oscillatory activity in frontal and auditory cortices

To study the characteristics of delta- and gamma-band rhythmic activity in frontal and auditory areas, we performed a cycle-by-cycle analysis of the recorded LFP. Cycles were considered only if they were part of consistent oscillatory activity (i.e. they were associated with a putative oscillatory bursts). Bursts of oscillatory activity were detected using the *bycycle* algorithm (Cole and Voytek, 2019), which capitalizes on a time-domain approach for quantifying waveform shape (**Fig. 2A**). A burst is detected based on four parameters, which control for signal-to-noise ratio (SNR) and waveform consistency (see Methods). In this context, an oscillatory burst occurs if the threshold values of these parameters are exceeded for at least 3 consecutive cycles.

**Figure 2.**
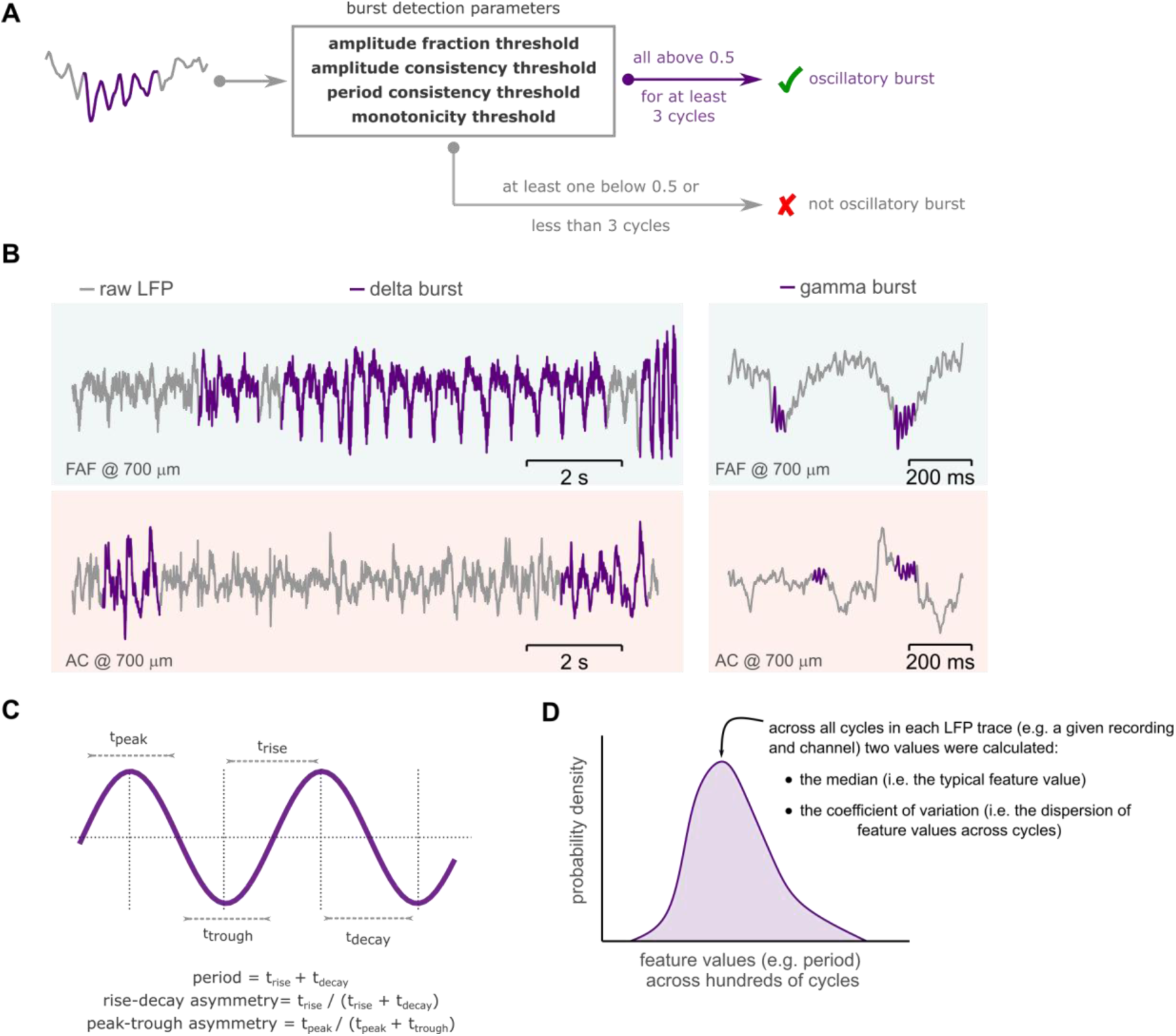
Burst cycle features and the coefficient of variation. (**A**) Schematic representation of the oscillatory burst detection algorithm. If at least 3 consecutive cycles fulfilled the detection parameters (enclosed in the box), these cycles together were considered as an oscillatory burst (marked in purple); otherwise, no burst was detected. No-burst cycles were not used in further analyses. (**B**) Representative delta- and gamma-frequency bursting activity (bursts marked in purple) in the FAF and AC, at a cortical depth of 700 μm. (**C**) Illustration of cycle waveform features: period, rise-decay asymmetry, and peak-trough asymmetry. An artificial sinusoidal waveform was utilized for illustrative purposes. (**D**) The median value for a given feature (e.g. period) across all cycles was used as the value of that feature for a given LFP trace. The coefficient of variation across all feature values was used as a measure of dispersion. Shown in the figure (in purple) is a schematic distribution of feature values for a given LFP recording.

Representative burst events in delta- and gamma-bands are shown for FAF and AC in **Fig. 2B**. The waveform shape of oscillatory activity was quantified by measuring three main features on a cycle-by-cycle basis: cycle period, cycle rise-decay asymmetry, and cycle peak-trough asymmetry (**Fig. 2C**; Cole and Voytek (2019)). For each LFP trace, the median feature value across all detected cycles was considered the waveform shape feature for that trace (**Fig. 2D**). The median summarizes a distribution of feature values, yielding one value per LFP trace (that is, 29 values for each FAF and AC channel). The median feature value of asymmetry metrics was normalized to 0.5 to account for possible confounds related to signal polarity differences in FAF and AC (see Methods; (Schaworonkow and Voytek, 2021)). To determine how and to what extent oscillatory waveform shape differed between recording locations, we performed systematic channel-by-channel pairwise comparisons. Only values obtained from simultaneously recorded LFP traces were compared to one another by using paired statistical testing. Median values for each ∼20-minute LFP were quantified from hundreds of cycles. That is, for Bat-01, no less than 665 and 370 delta-band cycles were used from FAF and AC, respectively, while no less than 829 and 146 gamma-band cycles were used from the same regions. For Bat-02, at least 561 and 468 delta-band cycles were used from FAF and AC, while at least 717 and 241 gamma-band cycles were used from the same areas.

### Bursting dynamics in frontal and auditory cortices

The data shown in **Fig. 1** suggest that the signal-to-noise ratio (SNR) of oscillatory activity in delta- and gamma bands is higher in FAF than in AC. We quantified SNR independently for each bat in frontal and auditory regions on a channel-by-channel basis (N = 15 observations for each channel in FAF or AC for Bat-01, and N = 14 for Bat-02). Distribution of SNR values are shown in **Fig. 3B** (top) for delta frequencies and in **Fig. 3E** (top) for gamma frequencies. Distributions from Bat-01 are shown with positive probability densities, whereas data from Bat-02 are given with negative probability densities merely for illustrative purposes. Note that the colour of each distribution corresponds to a specific channel in the shank, located at a certain cortical depth (see **Fig. 3A**). Given that the patterns across animals were highly consistent, we compared SNR values across recording sites by pooling data from the two bats. Channel-by-channel statistical comparisons revealed significant differences in SNR across recording sites (FDR-corrected Wilcoxon signed-rank tests, N = 29, significance when p_corr_ < 0.05). Comparisons are summarized in the matrices of **Fig. 3B-G** (bottom). A comparison matrix represents the effect sizes (d) of pairwise comparisons of SNR values across channels (|d| < 0.5 small, 0.5 <= |d| <= 0.8 medium, |d| > 0.8 large effect sizes; Cohen (2013)). A cell (*r, c*) in a matrix shows the effect size of comparing SNRs from a channel indexed by row *r*, and a channel indexed by column *c* (i.e. channel *r* vs. channel *c*). The relationship between a channel index and its relative depth in frontal or auditory cortex is schematized in **Fig. 3A** (notice the vertical lines next to channel numbers in **Fig. 3B-G** indicating cortical depths by following the colour schemes of **Fig. 1** and **Fig. 3A**). The upper right quadrant of each matrix represents comparisons of channels in FAF vs. those in AC. Only effect size values of significant comparisons (p_corr_ < 0.05) were shown; they were set to 0 otherwise. The matrices in **Fig. 3B** and **Fig. 3E** show strong differences in SNR between frontal and auditory cortices in the delta and gamma bands.

**Figure 3.**
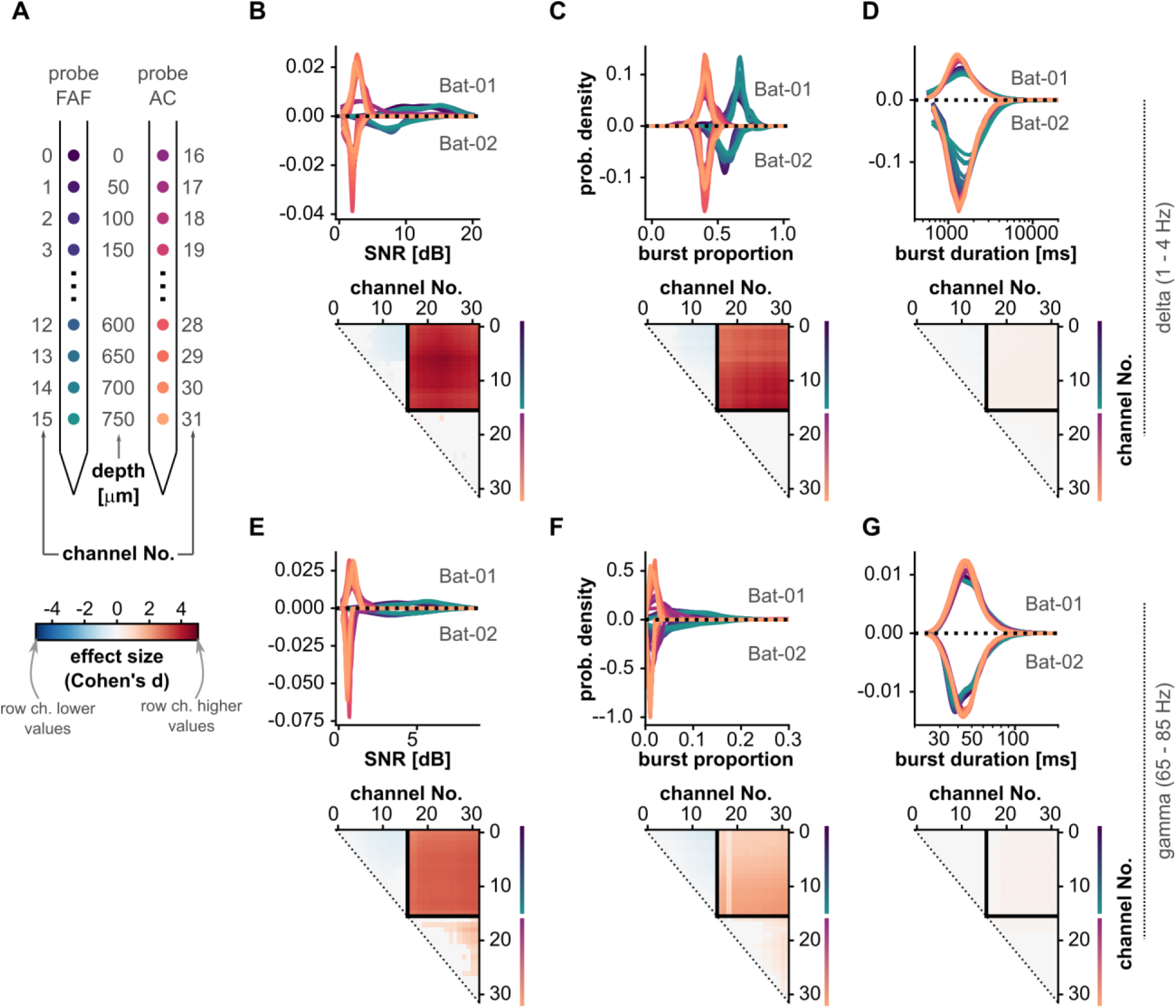
Bursting dynamics and signal-to-noise ratio in frontal and auditory cortices. (**A**) Schematic illustrating the relationship of region and cortical depth with the channel number markers of panels **B**-**G**. Notice that depths are colour-coded as in Fig.1. (**B**) *Top*: Distribution of signal-to-noise ratio (SNR) shown for each channel (notice colour schemes in panel **A** for the region and depth of each channel), across all recordings (N = 29 in FAF and AC). Values for Bat-01 are shown with densities > 0; values for Bat-02 are shown with densities < 0 only for illustrative purposes. *Bottom*: Effect sizes of channel-by-channel, pairwise statistical comparisons of SNR values (FDR-corrected Wilcoxon signed-rank tests). Effect sizes for comparisons that did not yield significance (i.e. pcorr >= 0.05) were set to 0. A cell (*r*, *c*) in the effect size matrix indicates the effect size of the comparison between burst proportion values in channel *r* and channel *c* (as per panel **A**). The quadrant spanning rows [0– 15] and columns [16–31] illustrates effect sizes of comparisons between channels in FAF and AC. In this quadrant, red colours indicate higher proportion values in FAF. (**C**) Same as in **B**, but data shown corresponds to burst proportions across recordings. (**D**) Same as in **C**, but data shown correspond to burst durations (note the logarithmic scale of the x-axis).

Typically, SNR values are interpreted solely on the basis of signal amplitude. However, high SNRs derived from the spectral properties of a signal could also indicate, beyond amplitude, a relatively high proportion of oscillatory events. We calculated bursting proportion as the ratio of the total time of bursting in an LFP trace relative to the total duration of that trace. Distribution of bursting proportions are given for both animals in **Fig. 3C** (top) for delta frequencies and in **Fig. 3F** (top) for gamma frequencies. From these data it appeared clear that, in both animals, the proportion of bursting events in FAF was higher than that in AC. Given this consistency, we pooled data across bats and compared on channel-by-channel basis bursting proportions across sites (FDR-corrected Wilcoxon signed-rank tests, N = 29, significance when p_corr_ < 0.05). These comparisons, summarized in the matrices of **Fig. 3C** (bottom) and **Fig. 3F** (bottom), corroborate that the proportion of delta- and gamma-band oscillatory events was significantly higher in FAF than in AC, with large effect sizes.

Higher proportion of bursting events could be influenced by two factors: more bursts occur in FAF than in AC, or bursts in FAF are longer than those in the auditory cortex (or both). To elucidate this, we examined the distributions of burst durations in delta- and gamma-band LFP traces from all channels. Distribution of burst durations are given independently for each animal in **Fig. 3D** (top) for delta and **Fig. 3G** (top) for gamma frequencies. To statistically compare burst durations, data across bats were pooled given the highly similar patterns observed from the two animals. For comparisons, all bursts from any given channel are considered, so the number of bursts per channel was not always the same (at least 2283 and 1977 bursts were used for delta frequencies from Bat-01 and Bat-02, respectively; in gamma, no less than 2984 and 1991 for each bat). Because of the uneven burst counts, channel-by-channel comparisons were not paired (FDR-corrected Wilcoxon ranksum tests, N >= 1991, significance when p_corr_ < 0.05). As readily visible from the distributions of burst duration, and as shown in the comparison matrices from **Fig. 3D** (bottom) and **Fig. 3G** (top), differences in burst durations between FAF and AC were statistically negligible, indicating that bursts in the FAF were more numerous, but not necessarily longer, than in the AC. This corresponds well with our initial observation of a very large burst density in frontal regions.

The data shown in **Figs. 1** and **3** demonstrate that spectral and bursting dynamics were highly consistent between Bat-01 and Bat02. Because of this consistency between animals (**Figs. 1** and **3**), data from the two bats were pooled in subsequent analyses.

### Bursting events in FAF and AC are temporally correlated

Transfer entropy analyses based on the phase of ongoing LFP activity, and direct electrical microstimulation of the frontal cortex to alter AC responsiveness, show that neural activity in FAF can significantly modulate its auditory cortical counterpart (Garcia-Rosales et al., 2022b). Given the functional and anatomical connections in the FAF-AC network, we sought to determine whether oscillatory bursts in one region are related to bursts occurring in the other. We reasoned that co-occurring bursts across brain structures could be a fingerprint of functional connectivity complementary to phase correlations (e.g. coherence), statistical dependencies (e.g. transfer entropy), or invasive approaches (e.g. electrical microstimulation). We calculated the burst co-occurrence index (**Fig. 4**), a metric that quantifies for any given channel the relationship between the onset of its own bursts with the occurrence of bursts in other channels (**Fig. 4A** illustrates how the index was computed). The burst co-occurrence index is shown in **Fig. 4B, C**, calculated for eight channels in total, four in each region, at representative depths of 50, 250, 450, 700 μm. Since the index is a cumulative count Z-normalized according to bootstrap distributions (see Methods), we could use it to evaluate the significance of burst co-occurrence across channels. Thus, red colours in **Fig. 4** indicate significant, temporally correlated increases in bursting activity in other channels (Z-values >= 6), while blue colours indicate significant, temporally correlated suppression of bursting activity in other cahnnels (Z-values <= -6). White colours indicate no significant deviations from baseline values.

**Figure 4.**
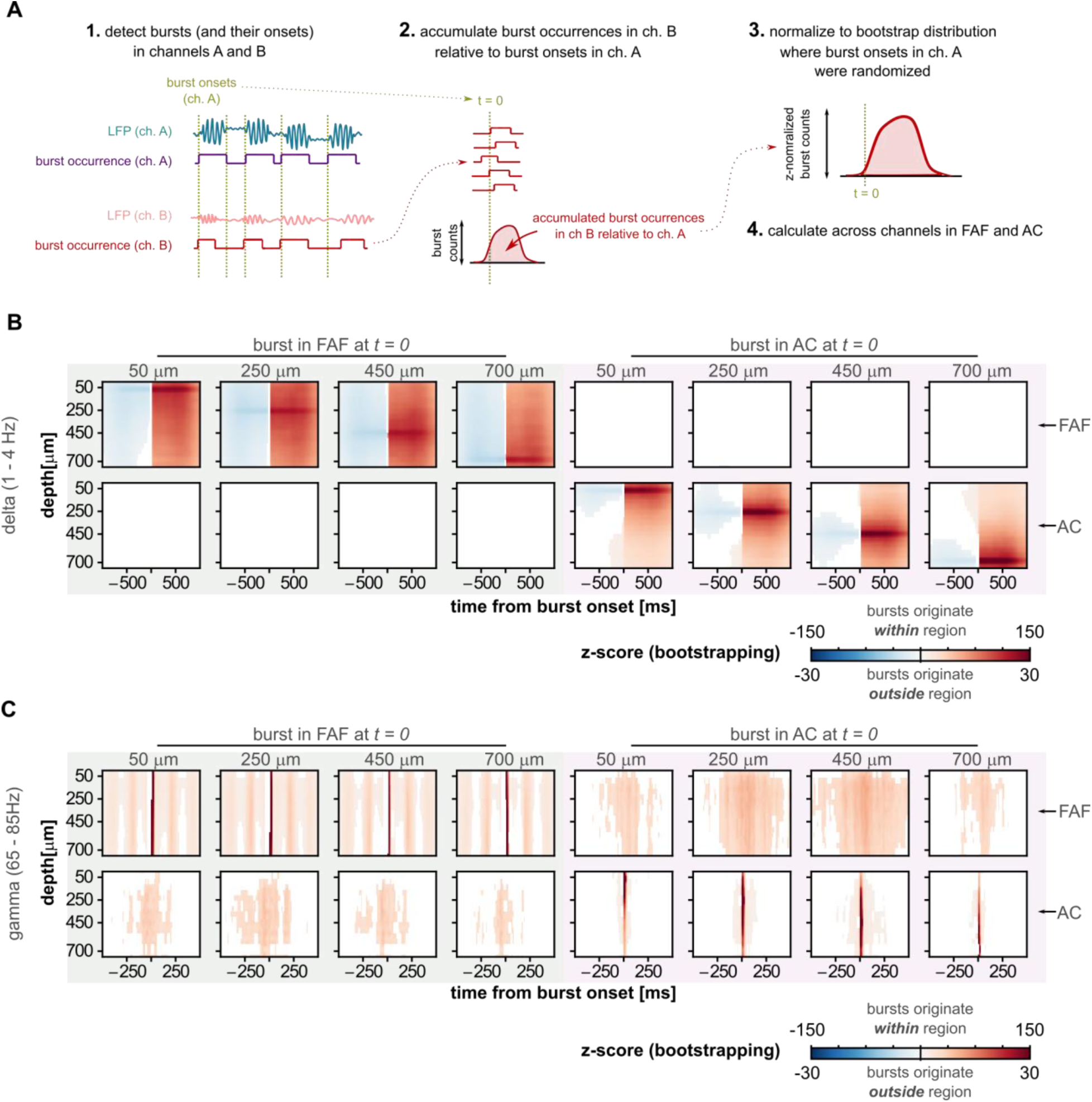
Burst co-occurrence in the FAF-AC circuit. (**A**) Schematic representation of the analysis for quantifying temporal co-occurrence of bursting events. Note that channels A and B could be drawn from the same or different cortical regions. (**B**) *Left* (shaded green): indicates delta-band burst co-occurrence across all channels, calculated relative to the onset of a burst at four representative depths (50, 250, 450, 700 μm) in the FAF. Matrices in the top row show burst co-occurrence in FAF channels; matrices in the bottom show co-occurrence of bursts in AC channels, aligned to burst onsets in FAF (at t = 0). Burst co-occurrence values were z-normalized relative to a bootstrapped baseline (blue colours: suppression of bursting activity; red colours: increased bursting activity). Only z-normalized values considered significant (|z| > 6, see Methods) are shown. *Right* (shaded pink): similar information, but in this case bursts originate in the AC at the same four representative depths (50, 250, 450, 700 μm). Here, t = 0 is aligned to auditory cortical burst onsets. (**C**) Same as in **B**, but depicting data related to the gamma frequency band. Note that burst co-occurrence matrices corresponding to bursts originating within a specific area (i.e. FAF or AC) are shown with a different colour scale than those corresponding to bursts originating outside a given area.

At delta frequencies (**Fig. 4B**), a burst onset in either FAF or AC was typically preceded by a suppression of bursting activity in channels of the same structure, and succeeded by a within-structure increase in burst co-occurrence across channels, peaking as trivially expected in the channel from which burst onsets were chosen. A similar pattern was observable for bursts detected in the gamma frequency range (**Fig. 4C**). However, we observed no clear pre-onset suppression in the gamma band, potentially due to much shorter durations of gamma bursts compared to delta ones. In gamma frequencies, an interesting pattern was evident: when burst onsets were taken from FAF channels (top left quadrant of **Fig. 4C**), a periodicity of burst co-occurrence emerged in the frontal area, with a temporal scale of ∼250 ms. This phenomenon constitutes evidence for strong coupling between gamma-band activity and low-frequency (delta) rhythms. These data resonate with that of a second study (and a different dataset) demonstrating clear coupling between the amplitude of gamma-band and the phase of delta-band LFPs in the FAF of *C. perspicillata* (Garcia-Rosales et al., 2022a).

Remarkably, when a burst onset occurred in frontal or auditory cortex, significant and consistent changes in burst co-occurrence in the other region happened for gamma-band LFPs (**Fig. 4C**). That is, bursts onsets in FAF were consistently and significantly correlated with gamma-band bursting in the AC, and vice-versa. Furthermore, **Fig. 4C** suggests a degree of spatial specificity to this relationship, wherein bursts originating in FAF appear more strongly related to those in middle layers of the AC (depths of 250-450 μm), while bursts originating in middle layers of the AC yield larger co-activation patterns in the FAF. Significant inter-areal burst co-occurrence was not equally clear in delta frequencies (**Fig. 4B**), although clear FAF-AC interactions occur in the delta band when considering transfer entropy analysis or even electrical stimulation experiments (Garcia-Rosales et al., 2022b). The apparent lack of burst interactions in the delta band shown in **Fig. 4B**, however, does not necessarily mean the absence of burst co-occurrence in these frequencies. Rather, this effect is a consequence of the stringency of the bootstrapping procedure (see Methods) interacting with the ubiquity of bursting activity in the FAF (**Fig. 3**). That is, bootstrap distributions were contaminated with real bursting activity when accumulating burst counts from the frontal cortex. Taken together, these results (particularly the ones related to gamma-band LFPs) suggest an intrinsic relationship between elevated bursting activity in frontal and auditory cortices, supporting the notion of strong functional connectivity in the FAF-AC network.

### Oscillatory waveform shape differences between frontal auditory cortices

We have shown the presence of oscillatory activity in delta and gamma frequencies in the frontal and auditory cortices of *C. perspicillata*. Oscillatory bursts across structures occur more often in the FAF, but are not necessarily longer than those in the AC. Remarkably, bursts in FAF and AC are temporally correlated, supporting the notion of concerted activity in the delta and gamma ranges in the FAF-AC circuit. Such correlated bursting activity occurs in very similar frequencies, yet they occur in functionally and anatomically distinct areas of the brain. Do these oscillations differ across structures?

A visual inspection of ongoing LFP activity revealed that the oscillatory waveform in the FAF was highly asymmetric (i.e. less sinusoidal, with more pronounced troughs), something that was not so obvious in the AC (see, for example, the representative bursts in **Fig. 2B**). The waveform shape of an oscillation was characterized by three main features (see above): period, rise-decay asymmetry, and peak-trough asymmetry. The distribution of feature values across recordings is given in **Fig. 5B-D** (top) for delta frequencies, and in **3E-G** (top) for gamma frequencies for all channels (see **Fig. 5A**; conventions are the same used for presenting data in **Fig. 3**). Note that the median feature value across all cycles is considered the feature value for a given LFP trace (**Fig. 2D**), thus yielding 29 feature values for each electrode either in FAF or AC (i.e. one value per recording). This allowed us to compare between recording sites using paired statistics, capitalizing on the fact that data in FAF and AC were simultaneously acquired.

**Figure 5.**
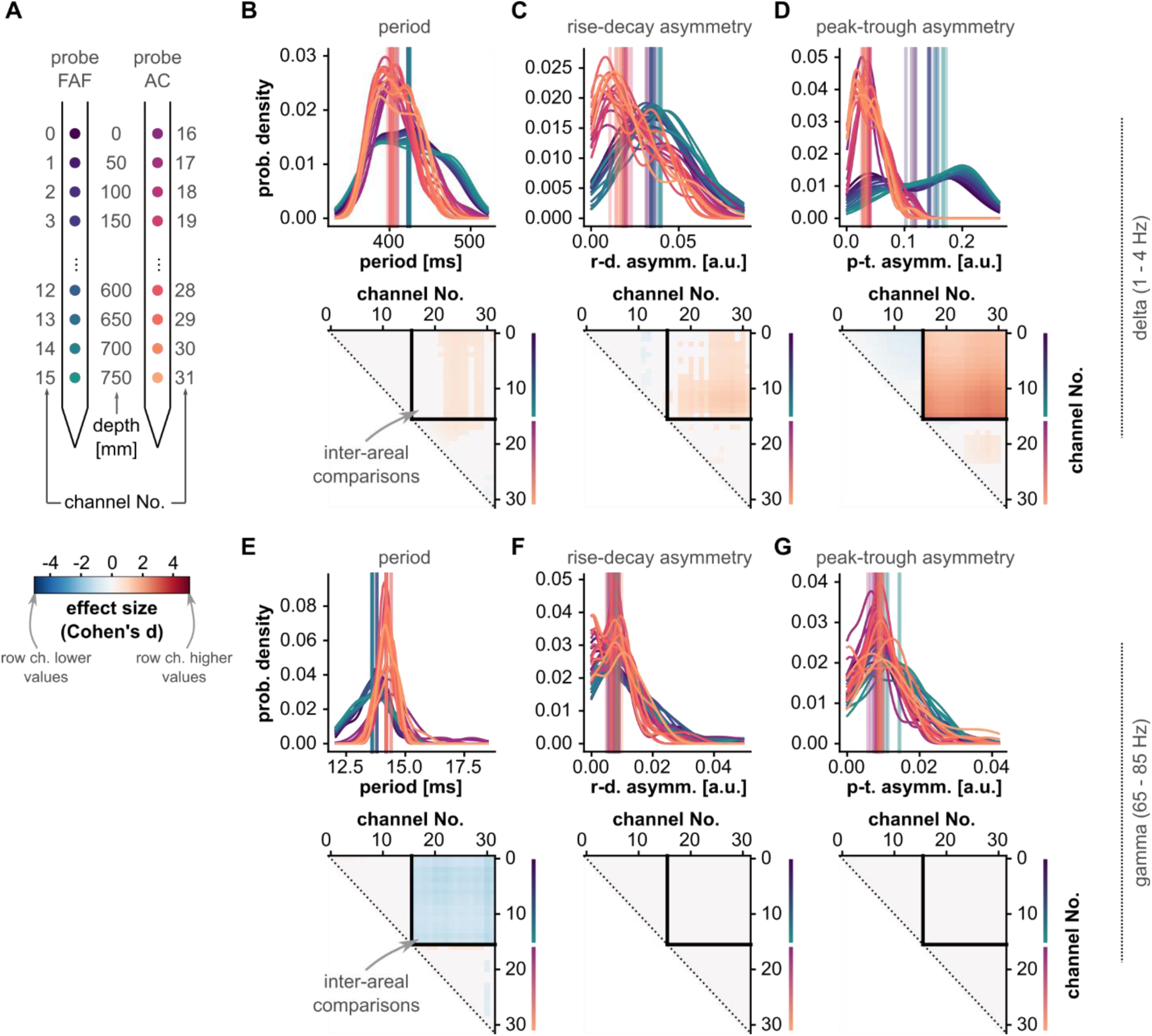
Waveform shape differences between frontal and auditory cortical LFPs. (**A**) Schematic illustrating the relationship of region and cortical depth with the channel number markers of panels **B**-**G**. Notice that depths are colour-coded as in Fig. 1. (**B**) *Top*: Distribution of oscillatory cycle periods across all recordings (N = 29; for each recording, the median period across all cycles is considered), for all channels (in FAF and AC; see panel **A** for region and depth according to colour), in the delta band. Vertical lines indicate the median of each distribution. *Bottom*: Effect sizes of pairwise statistical comparisons of population-level period values across all channels (FDR-corrected Wilcoxon signed-rank tests). Effect sizes for comparisons that did not yield significance (i.e. pcorr >= 0.05) were set to 0. A cell (*r, c*) in the effect size matrix indicates the effect size of the comparison between values in channel *r* and channel *c* (as per panel **A**). The quadrant spanning rows [0–15] and columns [16–31] illustrates effect sizes of comparisons between channels in FAF and AC. In this quadrant, blue colours indicate lower periods in FAF. (**C**) Same as in **B**, but corresponding to values of cycle feature “rise-decay asymmetry”. (**D**) Same as in **C**, but related to values of cycle feature “peak-trough asymmetry”. (**E**-**G**) Same as **B**-**D**, but shown for values obtained using gamma-band oscillatory cycles.

Channel-by-channel comparisons revealed significant differences across cortical regions (FDR-corrected Wilcoxon signed-rank tests, significance when p_corr_ < 0.05). These analyses are summarized in the comparison matrices of **Fig. 5B-G** (bottom; conventions are the same as those of **Fig. 3**). Delta-band oscillations in frontal and auditory cortices differed in period typically with small-to-medium effect sizes (|d| <= 0.8; **Fig. 5B**, bottom), but were strongly different in terms of their temporal asymmetries (**Fig. 5C-D**, bottom; |d| > 0.8 particularly for peak-trough asymmetries). The data in **Fig. 5** corroborates that the differences visible in **Fig. 2B** were consistent across recordings. Regarding gamma-band LFPs, the period of gamma-band cycles in FAF and AC differed more markedly than that of delta-band cycles (**Fig. 3E**, bottom; |d| > 0.8), although gamma-band oscillations differed only negligibly in their asymmetry across structures.

### Waveform shape variability is higher in auditory than in frontal cortex

By examining recordings independently we observed that beyond direct differences in waveform shape features (or lack thereof), feature values across cycles were typically less variable in the FAF than in the AC. That is, the distribution of feature values (e.g. period) were typically narrower for LFPs recorded in the frontal cortex. To evaluate the extent of this effect, we quantified for each LFP trace the variability of waveform shape features as the coefficient of variation (CV; **Fig. 2D**), and compared it across recording sites. The CV is a measure of dispersion, in the sense that it measures the “broadness” of a distribution. Thus, larger CVs indicate that cycle features vary over a wider range of possible values, suggesting a higher variability in the oscillatory processes. As with the median, the CV summarizes a distribution, yielding one value per LFP trace (see above and **Fig. 2D**). The same cycles used to calculate median feature values were used to calculate CV values.

The distributions of CV values across cycle features for each channel are given in **Fig. 6B-D** (top) for delta frequencies, and **Fig. 6E-G** (top) for gamma frequencies. CV values appeared consistently lower for channels in FAF than for those in AC. This trend was confirmed by statistical, channel-by-channel pairwise comparisons (FDR-correct Wilcoxon signed-rank tests, N = 29, significance when p_corr_ < 0.05), summarized in comparison matrices similar to those of **Fig. 3**. Statistical comparisons between channels located in different regions (the upper right quadrants of the comparison matrices) yielded the highest effect sizes (typically |d| > 0.8, large). CV values were consistently and significantly lower in FAF channels than in AC channels, in delta- and gamma frequency bands, for all cycle features. Some significant within-area differences also occurred (e.g. deeper channels in FAF had higher CV values than more superficial ones), yet effect sizes were typically medium (0.5 < |d| < 0.8) or small (|d| < 0.5). Overall, these results indicate that, beyond first-order differences in waveform shape, oscillatory activity in the frontal cortex exhibits a higher degree of cycle-by-cycle regularity (i.e. lower variability over cycles) than that of the AC. Note that such differences in regularity between regions are unlikely to arise from differences in the bursting proportions across FAF and AC (**Fig. 3**). Although more bursts result in more cycles contributing to a distribution, cycle-by-cycle regularity is quantified here using hundreds (sometimes thousands) of cycles obtained from relatively long LFP traces (ca. 20 minutes). These are well-sampled distributions whose CV should not be strongly affected by increasing the number of waveform shape features in them.

**Figure 6.**
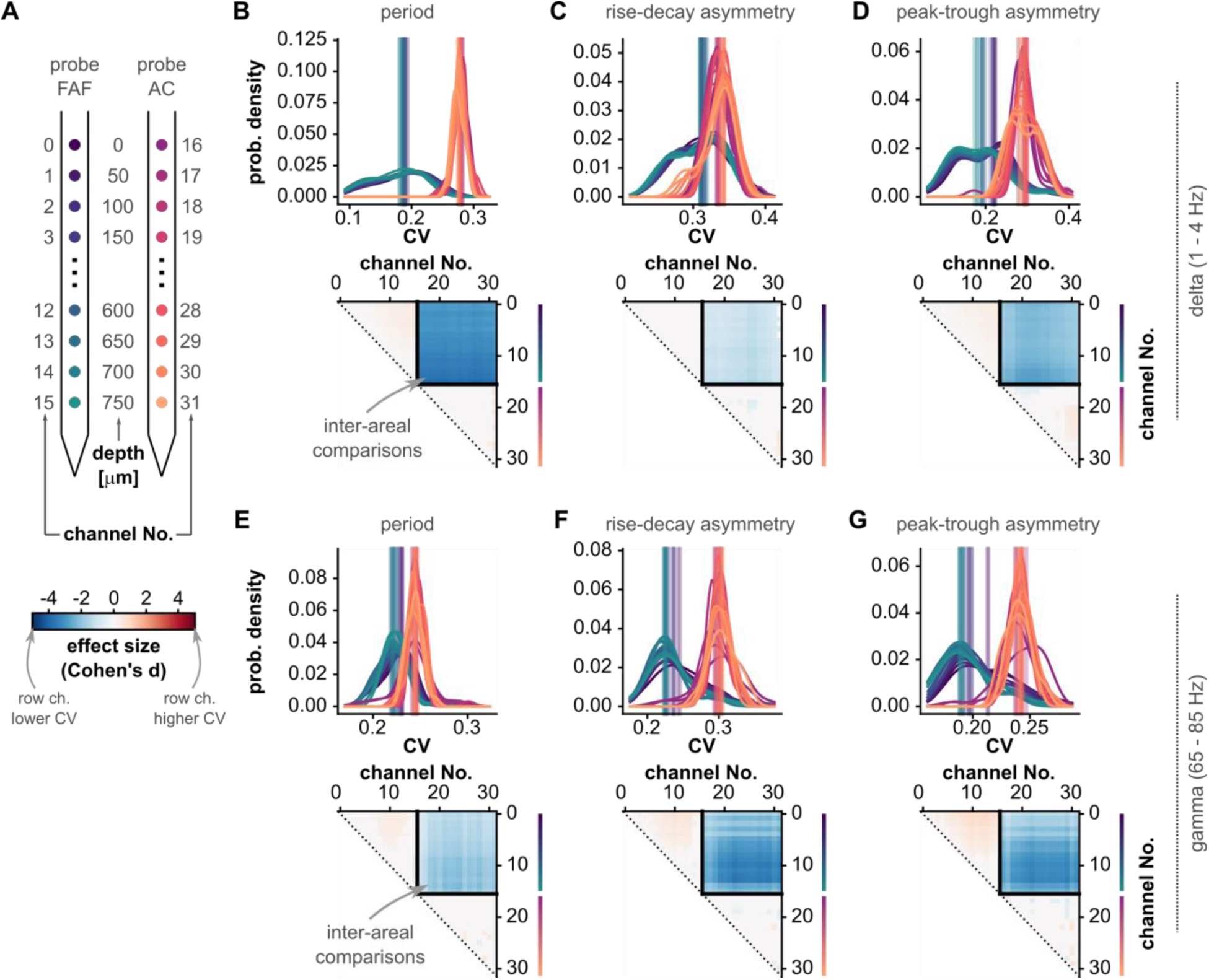
The variability of waveform shape features differs between frontal and auditory regions. (**A**) Schematic illustrating the relationship of region and cortical depth with the channel number markers of panels **B**-**G**. Notice that depths are colour-coded as in Fig.1. (**b**) *Top*: Distribution of CV values for oscillatory cycle periods across all recordings (N = 29), for all channels (in FAF and AC; see panel **A** for region and depth according to colour), in the delta band. Vertical lines indicate the median of each distribution. *Bottom*: Effect sizes of pairwise statistical comparisons of population CV values across all channels (FDR-corrected Wilcoxon signed-rank tests). Effect sizes for comparisons that did not yield significance (i.e. pcorr >= 0.05) were set to 0. A cell (*r, c*) in the effect size matrix indicates the effect size of the comparison between CV values in channel *r* and channel *c* (as per panel **A**). The quadrant spanning rows [0–15] and columns [16–31] illustrates effect sizes of comparisons between channels in FAF and AC. In this quadrant, blue colours indicate lower CV values in FAF. (**C**) Same as in **B**, but corresponding to CV values of cycle feature “rise-decay asymmetry”. (**D**) Same as in **C**, but related to CV values of cycle feature “peak-trough asymmetry”. (**E**-**G**) Same as **B**-**D**, but shown for CV values obtained using gamma-band oscillatory cycles. (Effect sizes can be interpreted as follows: |d| < 0.5 small, 0.5 <= |d| <= 0.8 medium, |d| > 0.8 large).

### Differences across regions are robust against burst detection parameters

The data indicate that oscillations in the FAF are more regular than those in the AC. However, the measurements of waveform shape used here can be affected by the SNR of the oscillatory activity used to quantify them. In particular, higher SNR of oscillatory activity in FAF (**Fig. 3B**) could result in narrower distributions of cycle features, because low SNR increases the variability of waveform shape features (see Schaworonkow and Nikulin (2019)). The SNR for burst detection is controlled by the parameter amplitude fraction threshold, which discards cycles below a certain amplitude percentile calculated from all cycles in an LFP trace (Cole and Voytek, 2019; Schaworonkow and Voytek, 2021). Therefore, to test whether the results shown above can be simply accounted for by different SNR levels in FAF and AC, we evaluated the sensitivity of the inter-areal differences to different values of amplitude fraction threshold in each region (**Fig. 7**).

**Figure 7.**
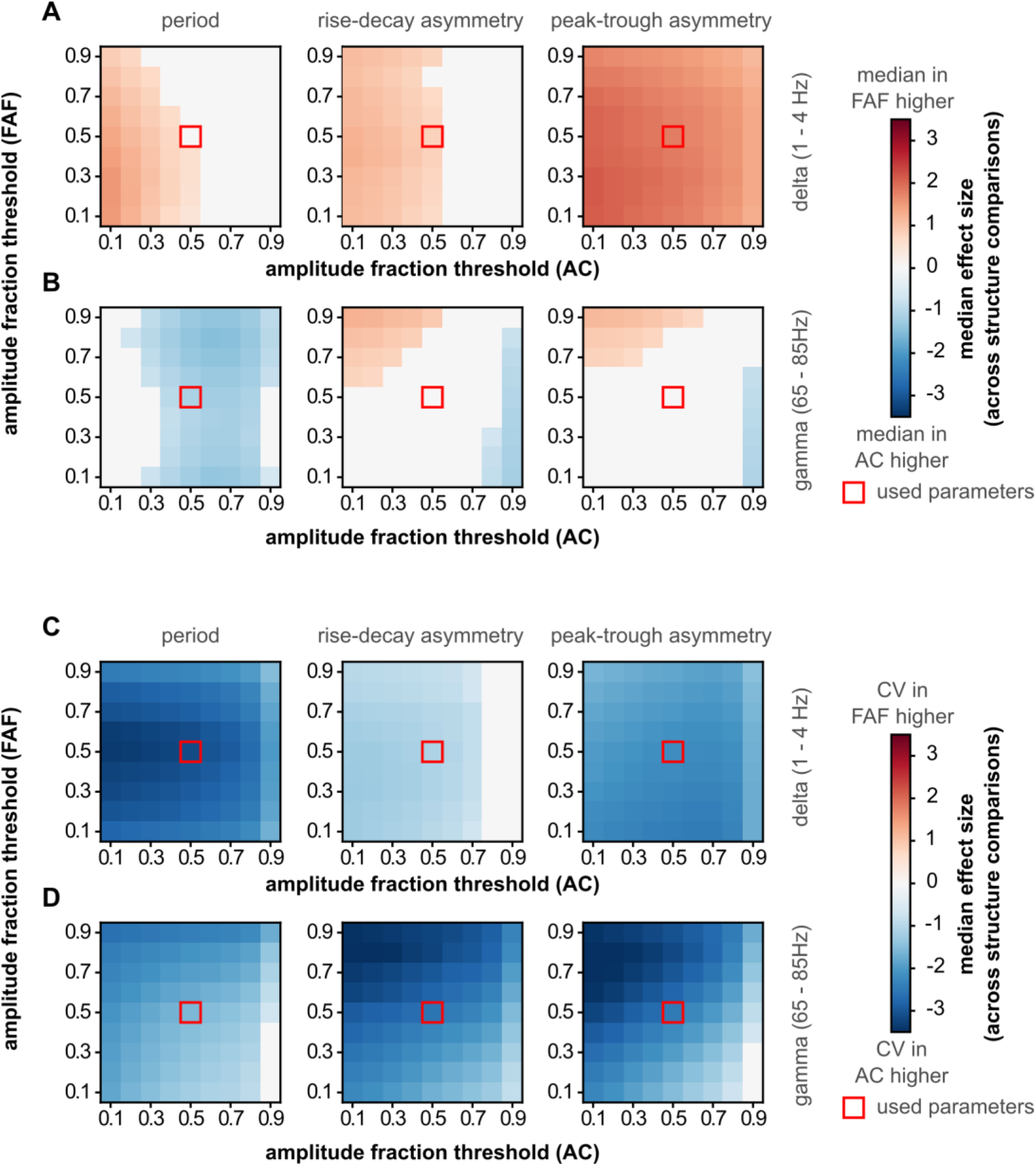
Differences in waveform shape features and their variability are robust against burst detection amplitude threshold. The burst detection parameter “amplitude fraction threshold” was varied independently in FAF and AC to determine whether SNR critically contributes to differences in oscillatory regularity between frontal and auditory areas. The difference across regions was measured as the median effect size obtained from comparing all pairs of channels in FAF and AC (e.g. median of the upper right quadrant in the comparison matrices in Fig. 5, labelled “inter-areal comparisons”). In the absence of significant differences (FDR-corrected Wilcoxon signed rank tests, pcorr < 0.05), effect size values were set to 0 (pcorr >= 0.05). (**A**) Median effect sizes across all values of amplitude fraction threshold tested in FAF and AC, for delta frequencies, comparing the median of cycle periods (left), cycle rise-decay asymmetries (middle) and cycle peak-trough asymmetries (right). (**B**) Same as in **A**, but data corresponds to cycles from gamma-band oscillatory bursts. (**C**, **D**) Same as in **a**, **b**, but the CV was calculated across cycle periods. Red squares indicate the amplitude fraction threshold values used to detect bursts used in the main results.

The median effect size of inter-areal comparisons was used as a summary metric of differences in median feature values (**Fig. 7A, B**) and CV values (**Fig. 7C, D**) across cortical regions. This metric corresponds to the median value of the upper-right quadrant of the comparison matrices in **Figs. 5** and **6**. We systematically varied the amplitude fraction threshold parameter (range: 0.1 – 0.9, step of 0.1) used to detect oscillatory bursts independently in the FAF or the AC, and for each iteration we calculated the median effect size of inter-areal comparisons. As depicted in **Fig. 7A**, period values for delta-band cycles were different between FAF and AC with typically medium or even low effect sizes (|d| < 0.5 for low, 0.5 <= |d| < 0.8 for medium), while asymmetries differed with typically strong effect sizes (|d| > 0.8) particularly when considering the peak-trough asymmetry as in **Fig. 5**. In general, observations across a broad range of threshold values conformed well to the data depicted in **Fig. 5** in delta- and gamma-bands (**Fig. 7B** for gamma). Those data were obtained with a threshold value of 0.5 (red squares in **Fig. 7**). Similarly, CVs were consistently lower in FAF than in AC across a wide range of amplitude fraction threshold values in both delta- and gamma frequencies (**Fig. 7C**, delta; **Fig. 7D**, gamma), for all three cycle features considered. These data were highly consistent with those shown in **Fig. 6**. These results indicate that the differences in waveform shape features and their CV values between frontal and auditory cortices are not trivially accounted for by differences in SNR across regions.

### A conceptual model captures patterns of waveform shape differences between FAF and AC

We hypothesized that differences across areas, particularly when considering the CV of waveform features, might reflect the activity of two distinct cortical generators exhibiting different degrees of regularity. We illustrate this idea with a conceptual model in which an oscillation occurs as a consequence of the temporally aligned rhythmic discharge of a population of neurons. This conceptualization makes no assumption on the nature of the neuronal oscillators themselves (see Discussion); instead, it only assumes that extracellular oscillatory activity occurs when a sufficiently large neuronal population fires concertedly (Buzsaki et al., 2012). We reasoned that a highly synchronous population firing would lead to a strong current at a specific phase of the LFP resulting in relatively asymmetric waveform shapes; by contrast, a relatively asynchronous population activity would yield less asymmetric temporal features. We simulated 30 neurons firing rhythmically for 300 seconds at a delta rate (3 Hz, for illustrative purposes; this can be generalized to other frequencies as well), with varying degrees of synchronicity among them. The synchronization of the spiking across neurons was manipulated by changing the duty cycle of a square pulse train determining to the instantaneous rate of an inhomogeneous Poisson process controlling a neuron’s firing rate (see Methods). Lower duty cycles represent narrower spiking windows and therefore higher synchronicity across neurons. From the neuronal firing in each condition, we generated a synthetic LFP by convoluting each spike train with a synaptic kernel and adding them over all neurons (**Fig. 8A**). This synthetic LFP was used to estimate cycle features computed with the *bycycle* algorithm, analogue to the analyses performed on the empirical data.

**Figure 8.**
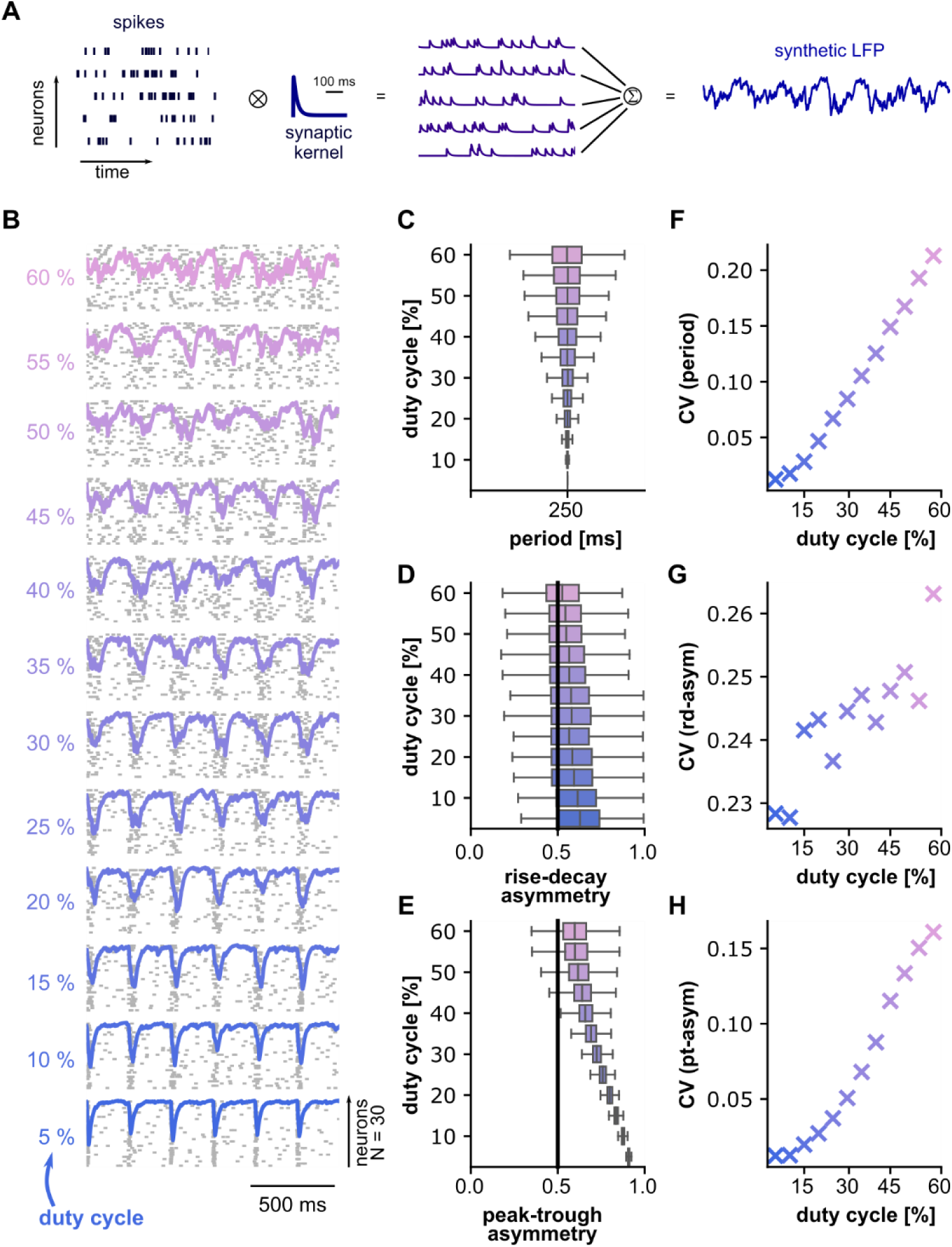
A linear model captures the differences in waveform shape between FAF and AC. (**A**) Schematic illustrating how synthetic LFP signals were derived from the spiking activity of a population of simulated neurons. (**B**) Representative spiking activity of a population of N=30 simulated neurons. The synchronicity across neurons varies with the duty cycle of a pulse train modulating firing rate (lower duty cycle, more synchronous; see Methods). A synthetic LFP was calculated for each condition (overlaid traces; see panel **A**). (**C-E**) The period (**C**), rise-decay asymmetry (**D**), and peak-trough asymmetry (**E**) values across all cycles detected by the *bycycle* algorithm for each duty cycle tested. The black line in panels **D** and **E** represents no asymmetry (i.e. a value of 0.5). (**F**-**H**) CV values of features period (**F**), rise-decay asymmetry (**G**), and peak-trough asymmetry (**H**). In panels **C**-**H**, values from each duty cycle simulation are colour coded according to panel **B**.

Simulated spiking activity with various degrees of synchronicity (controlled by the duty cycle parameter) is shown in **Fig. 8B** together with corresponding LFPs. Figure **8C-E** shows the distribution of cycle features (**Fig. 8C**, period; **Fig. 8D**, rise-decay asymmetry; **Fig. 8E**, peak-trough asymmetry) for each duty cycle condition. We did not observe changes in the median period across duty cycles. However, we did observe a consistent change in temporal asymmetries (**Fig. 8D, E**) indicating that a more synchronous neuronal population (lower duty cycles in the figure) resulted in more asymmetric waveform shapes. Note that the farther the median feature value is from 0.5 (black line in **Fig. 8D, E**) the more asymmetric LFP cycles are. In addition, we observed that as the neuronal population became less synchronized (i.e. higher duty cycles) feature values became more variable, as illustrated by the fact that the CV obtained from the feature distributions tended to increase together with the duty cycle (**Fig. 8G-H**). These two cases (higher asymmetry for more synchronized population spiking and more variability for less synchronized spiking) reflect differences in delta- and gamma-band oscillations in FAF and AC (i.e. higher asymmetry and less variability for oscillations in FAF), and offer a simple yet plausible account of the patterns observed across regions.

These results suggest that differences in FAF and AC waveform shape can at least be partially accounted for by different degrees of synchronicity in the underlying neuronal firing. To test this prediction, we turned to the spiking activity in frontal and auditory regions (**Fig. 9A**). Since oscillations were more asymmetric and less variable in FAF, we hypothesized that neuronal spiking would be more highly correlated in frontal than in auditory cortex and, additionally, more strongly synchronized with LFP oscillations (a secondary consequence of the model in **Fig. 8**). For each recording, we averaged correlation coefficients obtained from FAF and AC channels, and tested whether their values were significantly different across regions. These analyses corroborated that spike train correlations were higher in frontal regions (**Fig. 9B**, bottom; Wilcoxon signed-rank test, p = 2×10^-6^) with a large effect size (d = 0.84).

**Figure 9.**
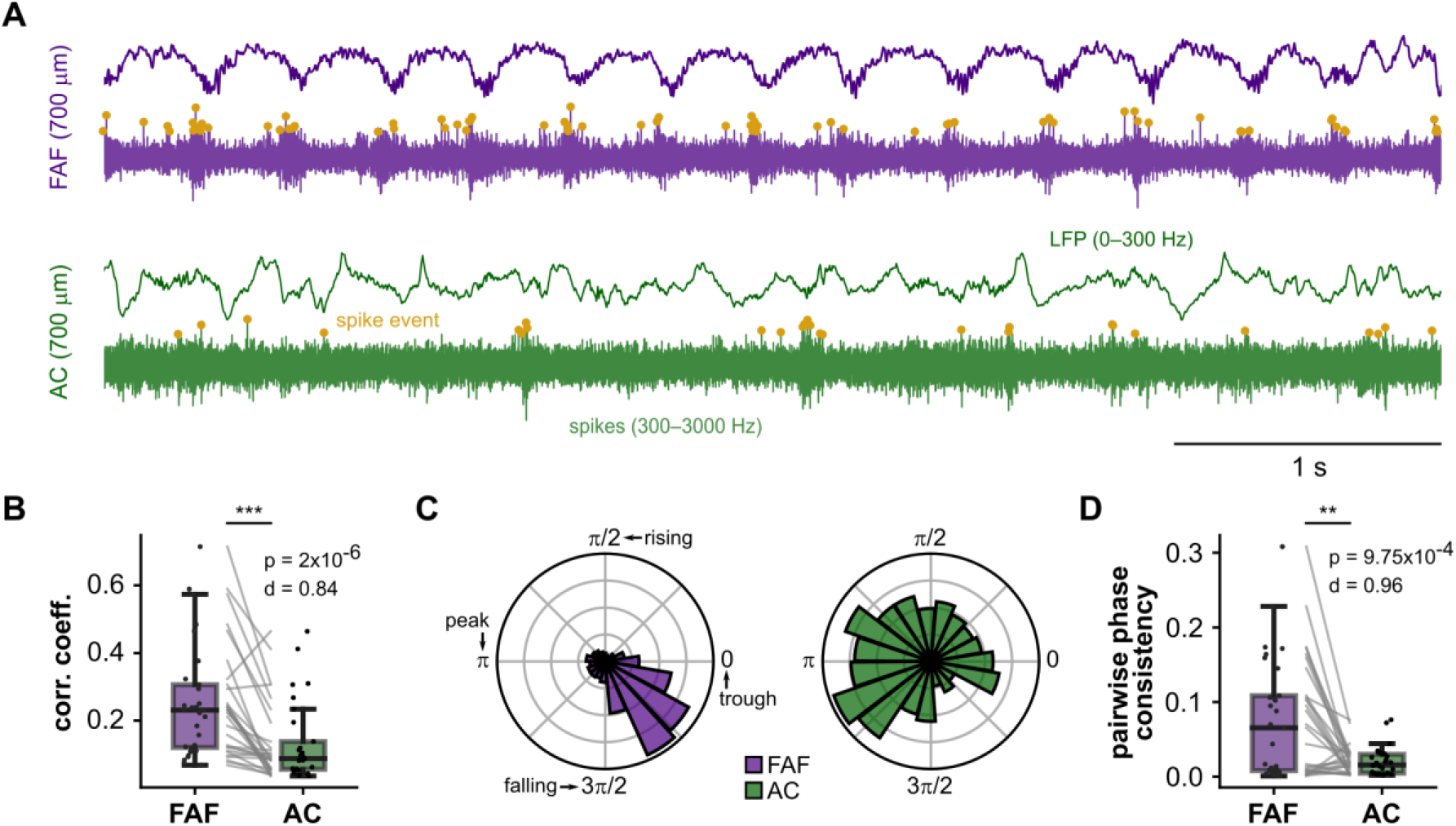
Spiking activity in FAF is more correlated and more strongly synchronized to delta-band oscillatory bursts. (**A**) Representative LFP (top) and spiking (bottom) activity from FAF (purple) and AC (green) electrodes at depths of 700 μm. (**B**) Spike-spike correlation coefficients for each recording in FAF and AC (N = 29; averaged across channels); spike-spike correlation in FAF was significantly larger than in AC (Wilcoxon signed-rank test, p=2×10^-6^, d = 0.84, large effect size). (**C**) Distribution of spike phases relative to delta LFPs in FAF (left, N = 6760 spikes) and AC (right, N = 661 spikes). Spikes were only those occurring during bursts of delta-band asctivity as detected by the *bycycle* algorithm. Troughs, peaks, rising and falling phases, for any given cycle, are indicated in the figure. (**D**) Average PPC values in FAF and AC were compared across all recordings (N = 29). There was significantly larger spike-phase consistency in FAF than in AC (Wilcoxon signed-rank test, p=9.75×10^-4^, d = 0.96, large effect size).

Because delta-band oscillations exhibited the largest differences in terms of asymmetry (**Fig. 5**), we studied spike-LFP relationships in this frequency range. Here, only spikes occurring within oscillatory bursts, as detected by the *bycycle* algorithm, were considered (note that these are the same bursts used in previous analyses). Spike times were expressed as the points of spike occurrence relative to the period of the burst cycle in which they occurred (0, spike occurs at beginning of cycle; 1, spike occurs at end of cycle), and spike phases were obtained by multiplying the relative spike timing by 2π. The distribution of spike phases from the recordings shown in **Fig. 9A** are depicted in **Fig. 9C** (N = 6760 spikes in FAF, N = 661 spikes in AC), suggesting a tighter clustering of spike phases in FAF. The pairwise phase consistency (PPC; Vinck et al. (2010)) was computed for all channels across recordings. The PPC measures how tightly spike phases group together (phase consistency) and constitutes a bias-free equivalent to the square of the phase locking value. Higher PPC values indicate higher spike-LFP coherence. To test whether spikes in FAF were more strongly synchronized to delta-band LFPs than those in the AC, we averaged PPC values across channels in FAF and AC (as described above) and statistically compared across regions. PPC values were significantly higher in FAF than in AC (**Fig. 9D**, bottom; Wilcoxon signed-rank test, p = 9.75×10^-4^) with a large effect size (d = 0.96).

Altogether, these results show that differences in waveform asymmetries between FAF and AC in delta frequencies are accompanied by differences in spike correlations and spike-LFP synchronization between regions. These observations are in line with predictions derived from the conceptual model illustrated in **Fig. 8**, and support a relationship between waveform shape and spike synchronization. Direct correlations between, for example, peak-trough asymmetry and spike-train correlations were, although significant, relatively weak (FAF, p = 0.033, adjusted R^2^ = 0.13; AC, p = 0.009, adjusted R^2^ = 0.2), indicating that oscillatory waveform shape cannot be trivially explained by local spike synchronization alone.

## Discussion

In this work, oscillations in the bat frontal and auditory cortices were studied with respect to their waveform shape. We show that oscillations present in simultaneously recorded LFPs in the fronto-auditory circuit differ markedly in waveform shape and in the variability of waveform features across individual cycles. This heterogeneity is not trivially accounted for by different levels of SNR in frontal and auditory regions. A conceptual model suggests a relationship between the temporal organization of neuronal spiking and waveform shape asymmetry, with higher spike temporal correlations leading to more asymmetric waveforms. In line with the predictions of the model, we demonstrate that spike-spike and spike-LFP correlations differ significantly in the FAF-AC network.

The bat frontal and auditory cortices are two brain regions with distinct cytoarchitectonic patterns, which likely accounts for the differences observed in oscillatory waveform shape across areas. *C. perspicillata*’s AC is a primary sensory region with a well-defined, six-layered columnar structure and clear inter-laminar boundaries (see Garcia-Rosales et al. (2019) for histology), following a blueprint that is typical across mammalian species (Douglas and Martin, 2004; Linden and Schreiner, 2003; Mountcastle, 1997). By contrast, *C. perspicillata*’s FAF lacks clear boundaries between layers (see Garcia-Rosales et al. (2022b); Weineck et al. (2020)), mirroring instead the stereotypical agranular or slightly agranular architecture of the mammalian frontal cortex (Beul and Hilgetag, 2014; Camarda and Bonavita, 1985; Shepherd, 2009). Differences between the bat frontal and auditory regions likely extend to other cytoarchitectonic properties such as the distribution of cell-type density and overall cellular organization. Beyond anatomy, cortical cytoarchitecture plays a significant role in defining activity patterns and brain function. Indeed, the functional characteristics of a given region are well-related to its cytoarchitecture (Badre and D’Esposito, 2009; Pandya and Yeterian, 1996), which includes the nature of incoming and outgoing axonal connections (Hilgetag et al., 2019; Kritzer et al., 1992; Passingham et al., 2002), cell-type specific characteristics (e.g. density, morphology; Benavides-Piccione et al. (2002); Beul and Hilgetag (2014)), and laminar organization (Hooks et al., 2011). Local cytoarchitecture affects neuronal firing patterns, which are known to vary consistently across functionally and anatomically well-defined regions (Badre and D’Esposito, 2009; Mochizuki et al., 2016; Shinomoto et al., 2009). Anatomical differences between granular and agranular cortical areas also result in distinct intra- and inter-laminar connectivity patterns (Beul and Hilgetag, 2014; Shepherd, 2009), which may also affect the dynamics of the generators of cortical oscillatory activity. Together, local anatomy, spiking patterns, and connectivity influence mesoscopic measurements of activity such as LFPs or other signals recorded non-invasively (Buzsaki et al., 2012; Cole and Voytek, 2017).

Other than local cytoarchitecture, respiration can also affect both single-neuron and oscillatory activities (Tort et al., 2018). For example, respiratory rhythms in mice entrain single neuron spiking and local cortical oscillations particularly –but not only-in frontal regions, (Koszeghy et al., 2018; Tort et al., 2018) with measurable functional consequences (Bagur et al., 2021; Folschweiller and Sauer, 2023). Likewise, heart rate fluctuations are known to correlate with brain oscillations, particularly during sleep (Mara and Julian, 2018; Mikutta et al., 2022). Respiratory or cardiac rhythms were not measured in this study, but their potential effects cannot be directly ruled out given that typical values for *C. perspicillata* lie close to delta frequencies: respiration rate, ∼2.5–4.5 Hz; hear rate: ∼8.33 Hz. For example, it is possible that respiration influences the patterns of rhythmicity and asymmetry overserved in frontal areas by directly modulating the LFP, by synchronizing neuronal spiking (thereby altering the LFPs), or a combination of both. Future studies should clarify the roles –if any-of respiration or heart rate in modifying oscillatory waveform shape dynamics.

Delta-band oscillations differed markedly across regions in terms of their temporal asymmetries, something that did not occur consistently for gamma-band activity (**Fig. 5**). However, for both frequency ranges we observed large and consistent inter-areal differences in the variability of shape feature values across individual cycles (**Fig. 6**). A conceptual model (**Fig. 8**) suggests that temporal asymmetries (i.e. waveform shape features) and their variability across cycles, could depend on the degree of correlated activity of the underlying neuronal population. The model in **Fig. 8** suggests that more synchronous populations yield highly asymmetric waveform shape and lower cycle-by-cycle variability, while less synchronous populations yield gradually a more sinusoidal shape with more variable cycle-by-cycle features. That waveforms become less asymmetric can be explained by temporal averaging of the contribution of each spike to the LFP, akin to the expected effects of spatial averaging in electro- or magneto-encephalographic recordings (Schaworonkow and Nikulin, 2019). Note that the model does not make any assumptions about important features of the underlying generators, such as location in the local circuitry, connectivity patterns, or component cell-types. As discussed above, these factors can influence both waveform shape and spiking dynamics. Instead, the model provides a parsimonious account of the empirical data shown in **Fig. 5**, assuming only that spiking is an important contributor to the LFP (Buzsaki et al., 2012). The model in **Fig. 8**, together with the waveform shape differences across regions, affords one prediction, namely that spike-spike and spike-LFP correlations should be higher in the area with more asymmetric signals (i.e. the FAF). Our results in **Fig. 9** corroborate such prediction, illustrating that the bat frontal cortex exhibits more correlated spiking, which is also more strongly synchronized the ongoing LFP phase in delta frequencies. As a concept, and supported by our data, the model draws a relationship between waveform shape asymmetry and the temporal dynamics of neuronal spiking in the neocortex.

A hypothesis stemming from the above observations is that differences in the variability of cycle features (measured by the CV) between FAF and AC might be explained by different values of temporal correlations in the underlying generators. In other words, it could be speculated that putative generators in the FAF operate with tighter parameters (reflected in higher temporal correlations) than their AC counterparts. One possible take on the functional implications of such phenomenon would be that frontal circuits rely more on internal timescales, while auditory circuits exhibit an elevated flexibility and perturbability. Previous studies have demonstrated that activity patterns in the rodent prefrontal cortex exhibit less variability than those of sensory regions (Castano-Prat et al., 2017; Ruiz-Mejias et al., 2011), potentially reflecting a cortical hierarchy of excitability and circuit properties. In such hierarchy, peripheral areas exhibit more adaptability to sensory stimuli (and therefore more variability), while frontal areas exhibit higher stimulus independence, yielding activity patterns better related to local network dynamics (Badre and D’Esposito, 2009; Braun and Mattia, 2010; Ruiz-Mejias et al., 2011)). In the bat brain, the FAF appears to be a modulation and control structure that may also be involved in the integration of diverse inputs during echolocation and navigation, as reflected by its internal dynamics and by the anatomical and functional connectivity patterns with other cortical and subcortical regions (Casseday et al., 1989; Eiermann and Esser, 2000; Garcia-Rosales et al., 2022b; Kanwal et al., 2000; Kobler et al., 1987; Weineck et al., 2020). Conversely, the bat AC (as that of other mammals) is primarily tasked with representing sounds that may unfold in time over nested timescales, typically exhibiting varying degrees of periodicity which require higher adaptability and flexibility (Doelling et al., 2019; Garcia-Rosales et al., 2018; Henry and Obleser, 2012; Lakatos et al., 2013; Teng et al., 2017). Indeed, previous modelling work suggests that neuronal response patterns in FAF and AC can be accounted for by slower synaptic dynamics in the frontal region (Lopez-Jury et al., 2020), something that could be detrimental for precise stimulus tracking but that could be important for sensory integration. From the above, we hypothesize that a higher level of variability in the auditory cortical circuitry (**Fig. 6**) might aid with efficient sensory representations in AC (see Pittman-Polletta et al. (2021)), while narrower dynamics could be important for high-level computations in FAF (e.g. sensory integration), closely tied to internal timescales and more robust against external perturbations.

In conclusion, we have shown that simultaneously recorded oscillatory activity across frontal and auditory cortices differs markedly in waveform shape. Additionally, a conceptual model, paired with empirical results, suggests a relationship between waveform shape and local spiking activity. This intriguing relationship could serve as a tool for constraining generative models of neural oscillations, and can be used to draw hypotheses after observing waveform shape differences across experimental conditions. The oscillations studied here in frontal and auditory regions occur in similar frequencies and are functionally related (**Fig. 4**; (Garcia-Rosales et al., 2022b)), but they nevertheless possess distinct dynamics that reflect the heterogeneous anatomical and functional properties of the bat fronto-auditory network.

## Acknowledgments

This work was supported by the DFG (Grant No. HE 7478/1-1, to J.C.H.), the Joachim-Herz Foundation (fellowship granted to F.G.R.), and a Marie Skłodowska-Curie Actions Postdoctoral Fellowship (grant No. 101062497, to N.S.).

## Author Contributions

F.G.R, N.S., and J.C.H. conceived and designed the research. F.G.R collected and analysed the data, produced original figures, and wrote the first draft of the manuscript. F.G.R., N.S, and J.C.H. discussed analyses and results, interpreted data, and reviewed figures and text.

